# Zebrafish use visual cues and geometric relationships to form a spatial memory

**DOI:** 10.1101/620575

**Authors:** Ksenia Yashina, Álvaro Tejero-Cantero, Andreas Herz, Herwig Baier

## Abstract

Animals use salient cues to navigate in their environment, but their specific cognitive strategies are largely unknown. We developed a conditioned place avoidance paradigm to discover whether and how zebrafish form spatial memories in a Y-shaped maze. Juvenile zebrafish, older than three weeks, learned to avoid the arm of the maze that was cued with a mild electric shock. We found that the fish required distinct visual patterns to develop a conditioned response. Interestingly, individual fish solve this task in different ways: by staying in the safe center of the maze, by preference for one, or both, of the safe patterns, or by mixed strategies. In experiments in which the learned patterns were swapped, rotated or replaced, the animals could transfer the association of safety to a different arm or to a different pattern using either visual cues or location as the conditioned stimulus. These findings show that juvenile zebrafish exhibit several complementary spatial learning modes and pave the way for neurobiological studies of navigational mechanisms in this model species.

## Introduction

When performing a behaviorally relevant task, such as seeking a refuge or a feeding spot, animals rely on various environmental cues to memorize and recall specific locations (Cheng, 1986; Durán, Ocaña, Broglio, Rodríguez, & Salas, 2010; Eichenbaum, 2017; Franz & Mallot, 2000; Gouteux, Thinus-Blanc, & Vauclair, 2001; Kelly, Spetch, & Heth, 1998; Salas et al., 2006; Vallortigara, Zanforlin, & Pasti, 1990). Teleost species, such as adult goldfish, have been shown to use several strategies to navigate in diverse locales, and may associate a reward with specific places, with geometric configurations of the environment, or with local visual cues (López, Bingman, Rodríguez, Gómez, & Salas, 2000; López, Broglio, Rodríguez, Thinus-Blanc, & Salas, 1999; Vargas, López, Salas, & Thinus-Blanc, 2004). The reward can also be associated with a combination of these cues, e.g., with the animal’s location in a plus-shaped maze and a salient visual stimulus. When the visual cue is relocated, the cue-guided strategy becomes incompatible with a location-guided strategy, and the fish may choose one of the two strategies. Surprisingly, some fish choose a strategy of following the visual cue, while others use a strategy of seeking the location (López et al., 2000).

Few studies have addressed the question of spatial learning strategies in zebrafish – a genetically tractable model organism. Adult zebrafish are capable of associative learning using, among others, olfactory, social, and place cues (Al-Imari & Gerlai, 2008; Aoki, Tsuboi, & Okamoto, 2015; Braubach, Wood, Gadbois, Fine, & Croll, 2009; Kalueff et al., 2013; Kenney, Scott, Josselyn, & Frankland, 2017; Lal et al., 2018). Much less is known about the learning abilities of larval and juvenile zebrafish before the age of four weeks, a period during which the brain is accessible to non-invasive imaging approaches. There have been reports of learning effects in one-week-old larval zebrafish using classical conditioning paradigms (Aizenberg & Schuman, 2011; Harmon, Magaram, McLean, & Raman, 2017; Lee et al., 2010) and operant conditioning paradigms (Hinz, Aizenberg, Tushev, & Schuman, 2013; Yang, Meng, Li, & Wen, 2019). Robust learning effects were observed in three-week-old juvenile zebrafish (Matsuda, Yoshida, Kawakami, Hibi, & Shimizu, 2017; Valente, Huang, Portugues, & Engert, 2012). While these studies demonstrated the ability of young fish to associate cues with a reward or a punishment, the spatial component of learning has not yet been explored.

To investigate the mechanisms underlying the flexibility in selecting specific navigation strategies, we developed a Y-maze paradigm. Here, fish were conditioned to avoid one of three arms of the Y-maze by cueing one arm with electric shocks. This experimental setup allows fish to explore several compartments of the maze in an operant mode, while offering the experimenter full control of the fish’s visual environment. We characterized the learning behavior of zebrafish in the Y-maze, and found robust learning effects only in juvenile animals older than three weeks. Experiments in which we replaced, swapped or rotated the visual patterns showed that the animals used a variety of strategies to memorize the safe areas in the maze. Some fish avoided both the conditioned arm and all other arms by staying in the center of the maze, while others preferred either a safe pattern or a safe location in the maze. These findings indicate that zebrafish use visual cues, in conjunction with geometric relationships, to navigate through their environment.

## Results

### Operant conditioning in a Y-maze as a readout of spatial memory

We configured a conditioned place avoidance paradigm (CPA), in which young zebrafish could explore a Y-maze. Each arm of the maze had a distinct visual pattern associated with it projected from below (Figure 1A). Experiments consisted of three consecutive sessions. In the first session (habituation) fish were free to explore the maze. At the end of this session we identified the arm with the highest occupancy as the preferred arm of each fish, which was then selected as the arm for conditioning. During the second session (conditioning) we trained the fish to avoid the conditioned arm by punishing entry into that arm with a mild electric shock. In the third session (test) the electric stimulation was switched off and the memory of the fish was tested.

**Figure 1.**
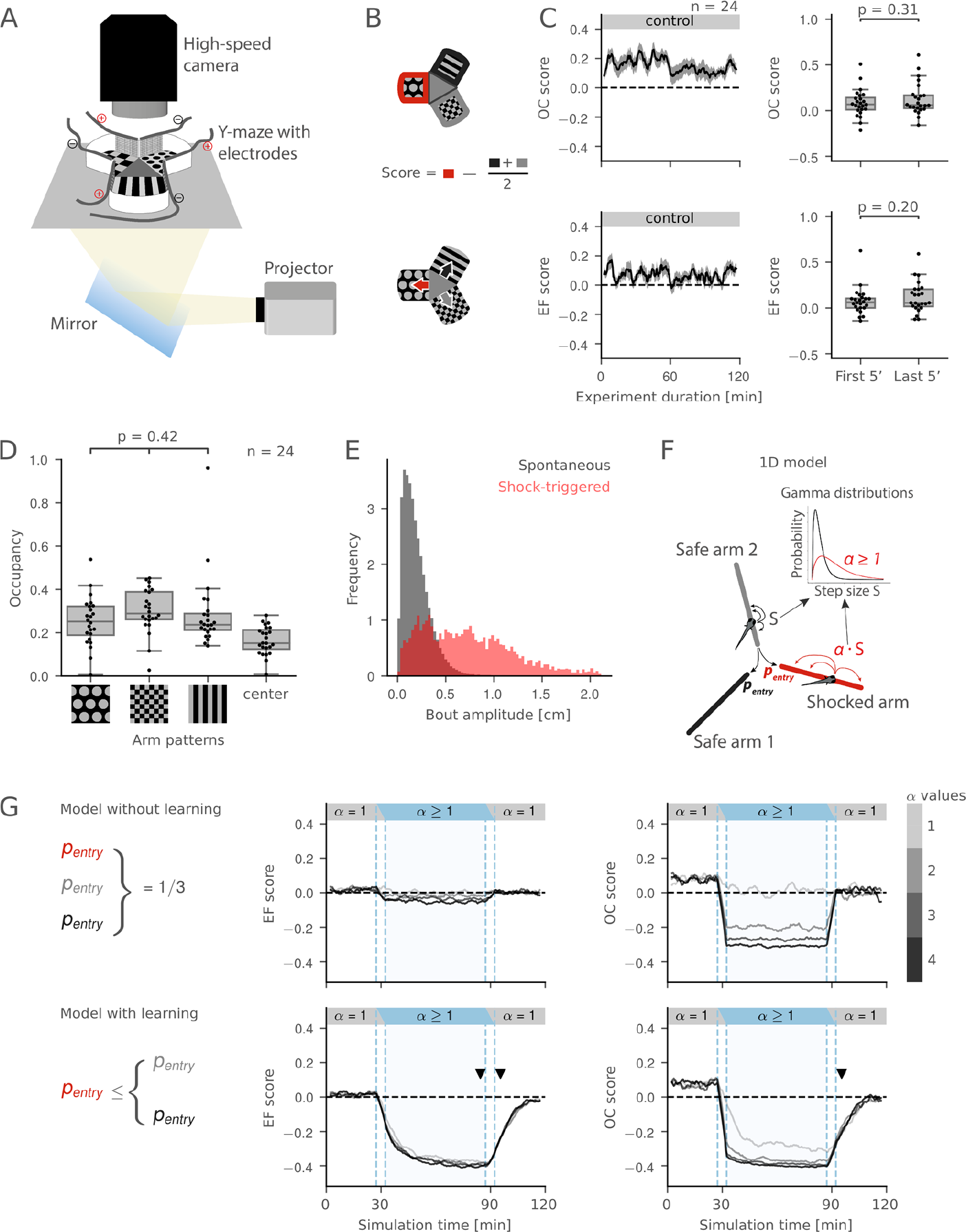
The conditioned place avoidance (CPA) paradigm. **(A)** Experimental setup. **(B)** Measures of fish performance in the paradigm, top: arm occupancy (OC); bottom: arm entry frequency (EF); middle: schematic for the calculation of the preference scores for OC/EF. Positive values of the scores correspond to arm preference, negative values correspond to arm avoidance. Red color represents the shocked arm, black and gray represent the safe arms. **(C)** Left: moving averages of the OC/EF scores in a 2-hour control experiment (mean ± s.e.m.). Right: box plots for OC/EF scores in the first and the last 5 minutes of the control experiments (Mann-Whitney test, n = 24 fish). Box plots show median and quartiles; whiskers show 1.5× interquartile range; dots show values for individual fish. **(D)** There is no significant preference for any of the visual patterns (ANOVA, n = 24 fish). **(E)** Distributions of amplitudes for spontaneous (gray) and shock-triggered swim bouts (red). **(F)** Pseudo-random 1D walk model used to evaluate CPA measures. Bout amplitude *S* for simulated movement was drawn from a Gamma distribution, amplitude in the conditioned arm was multiplied by a speed ratio *α*. **(G)** Top: model without learning. Bottom: model with learning. Learning rule was implemented by decreasing the probability of entry into the conditioned arm. Middle and right: moving averages of EF/OC scores. Black/gray lines correspond to simulations with different speed ratios *α*. Dashed vertical blue lines mark the areas where moving average combined information from two sessions. Arrowheads mark the differences in the OC/EF scores between top and bottom. Moving averages are calculated with a 5-min time window and a 30-s time step.

We developed two measures to estimate the effects of conditioning (Figure 1B). Our first measure was the occupancy (OC) of each arm, or center of the maze, which we calculated as the proportion of the total experiment time the fish spent in the respective part of the maze. Our second measure was the entry frequency (EF) of each arm, calculated as the number of entries into the respective arm divided by the total amount of arm entries. We did not calculate EF measure for the center of the maze, because it was equal to the sum of all arm entries and did not reflect avoidance/preference of maze arms. Using the OC measure, we checked if fish had any intrinsic preference for the visual patterns used in the experiments. We ran control experiments, in which the fish were allowed to swim in the maze for two hours without electrical stimulation. The fish did not show a preference for any of the visual patterns (Figure 1D).

For each measure (EF or OC) we created an associated score to evaluate the difference in avoidance/preference of the maze arms (the center of the maze was not included). The score was calculated by subtracting the average of the measures in the two safe arms from the measure in the conditioned arm (Figure 1B, see schematic equation). This score is positive when a fish prefers the conditioned arm and negative when the fish avoids it. We evaluated the stability of the OC and EF scores in the control experiments. The OC and EF scores of the arm that was preferred in the first 30 minutes of the experiment stayed positive on average over the duration of the control experiment (Figure 1C). This suggests that the arm preference is stable and any change in CPA measures in conditioning experiments is a result of conditioning and not of stochastic variation in the fish’s arm preferences.

### A biologically realistic agent model replicates learned avoidance behavior

Larval and juvenile zebrafish swim in characteristic swim bouts (Supplementary Video 1). Electric stimulation caused swim bouts of increased amplitude compared to the amplitude of spontaneous swim bouts (Figure 1E). We hypothesized that increased swimming speeds under electric stimulation in the conditioned arm could lead to changes in the OC and EF measures independent of the learning abilities of the fish. We developed a null-model of the CPA measures in the conditions of stimulation-enhanced swim bouts to test our hypothesis (Figure 1F, see Methods). In this computational model the simulated agent could move in pseudo-random walk fashion in a 1-dimensional arm in discrete bouts of size *S*, with a certain probability of moving into a different arm. The effect of electric shocks on the speed of the fish was simulated by multiplying the bout size in the conditioned arm by a factor *α* ≥ 1. Learning was excluded from the design of this basic model. We generated fish trajectories for different values of *α* and found that the OC score of the conditioned arm, but not its EF score, was dramatically decreased during the conditioning session (Figure 1G, top). This result highlights the effect of increased speed on arm occupancy that is independent of learning. We then added a learning rule to the model by decreasing the probability of entry into the conditioned arm, and observed a decrease in both the OC and EF scores of the conditioned arm in conditioning and test sessions (Figure 1G, bottom). We concluded that the decrease in the occupancy of the conditioned arm while the fish is shocked is not sufficient for inferring learning, as it can be explained by learning-unrelated reasons, and decided not to use the OC score during conditioning.

### Spatial learning abilities emerge at juvenile stages

We used the established paradigm to identify the earliest developmental stage at which zebrafish can be conditioned. We chose three age groups: 1-, 2-, and 3-week-old fish; and performed experiments that consisted of two sessions: habituation and conditioning (Figure 2A). We found that fish had an EF score significantly lower than zero at the end of conditioning only once they reached 3-weeks of age (Figure 2B). The OC score was significantly decreased in all age groups at the end of conditioning; however, this result was predicted by our computational model and could be unrelated to the learning ability of the animals (Figure 2C). The size of the fish was more variable at later stages of development, suggesting that fish of the same age could be at different developmental stages (Figure 2D).

**Figure 2.**
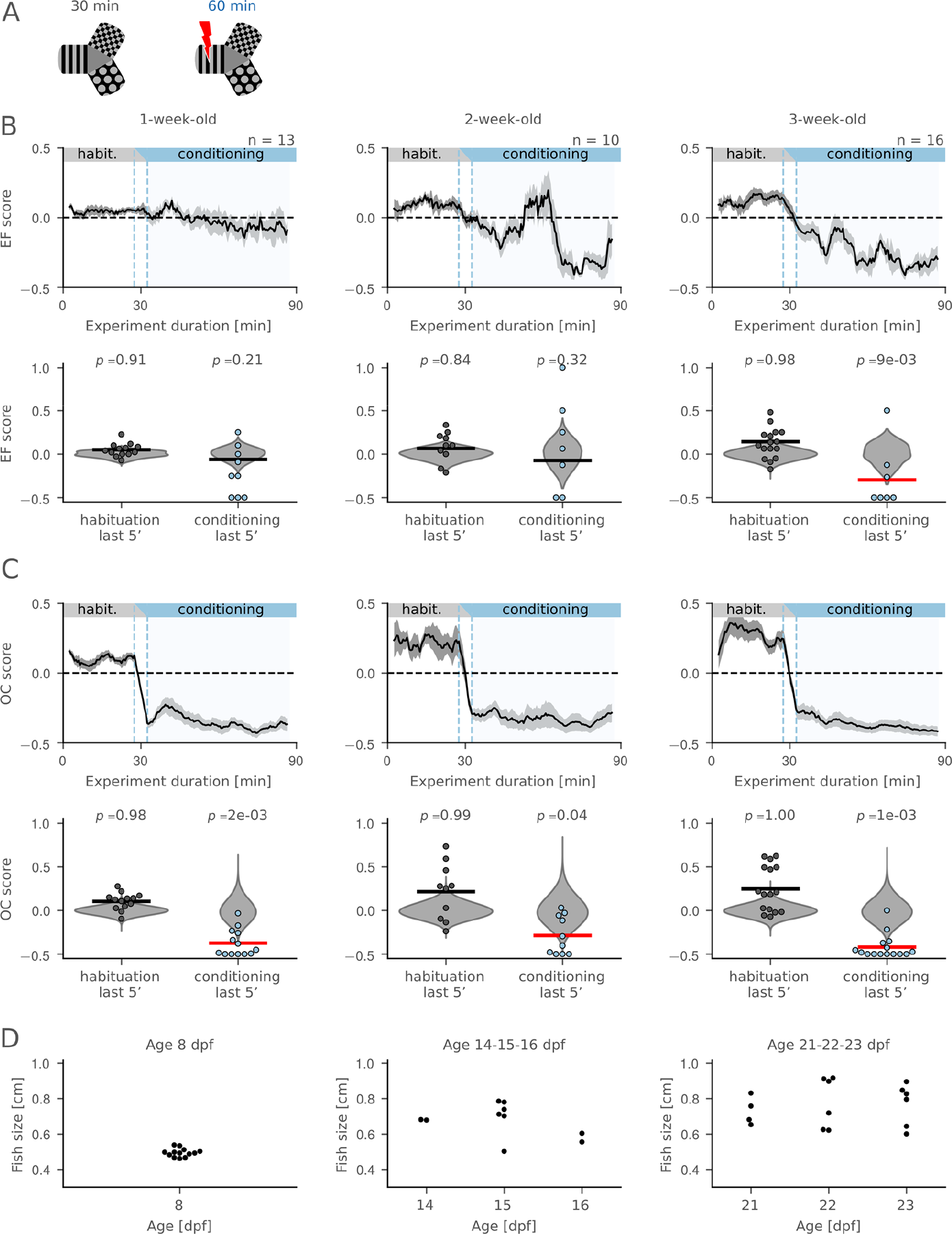
Performance in the CPA paradigm across different age groups. **(A)** Schematic of the protocol: habituation and conditioning sessions. **(B)** Changes in EF scores across different age groups. Note that the EF score becomes significantly lower than zero only in 3-week-old fish. Top: moving average (mean ± s.e.m.). Dashed vertical blue lines mark the areas where moving average combined information from two adjacent sessions. Bottom: comparison of the EF scores in the last 5 min of conditioning with the null-distribution (permutation test). Individual values for each fish are shown as dots, null-distributions are shown in gray violin plots. Horizontal lines show the sample means, the line is red if the mean lies to the left of the 5^th^ percentile in the null-distribution. **(C)** Changes in OC scores across different age groups. Figure annotations are the same as in (B). **(D)** Distribution of body sizes across different ages. Average body size and its variability increase with age: 1 week 4.95 ± 0.23 mm; 2 weeks 6.74 ± 0.93 mm; 3 weeks 7.64 ± 1.15 mm, mean ± SD. Dpf, days post fertilization.

Based on this comparison of different age groups we selected 3-week-old fish for further experiments.

### Memory of visually cued place persists for at least ten minutes

We tested the duration of the aversive memory formed at the end of conditioning in the third (test) session to the protocol, during which electric shocks were switched off (Figure 3A). Analysis of the OC score revealed that fish had formed a memory of the aversive arm: the OC score was below zero in the first 10 minutes of the test session (Figure 3A, permutation test *p*-value = 7⋅10^−4^, n = 40 fish). At the end of the test session the OC score returned to zero (Figure 3A, permutation test *p*-value = 0.21, n = 40 fish)

**Figure 3.**
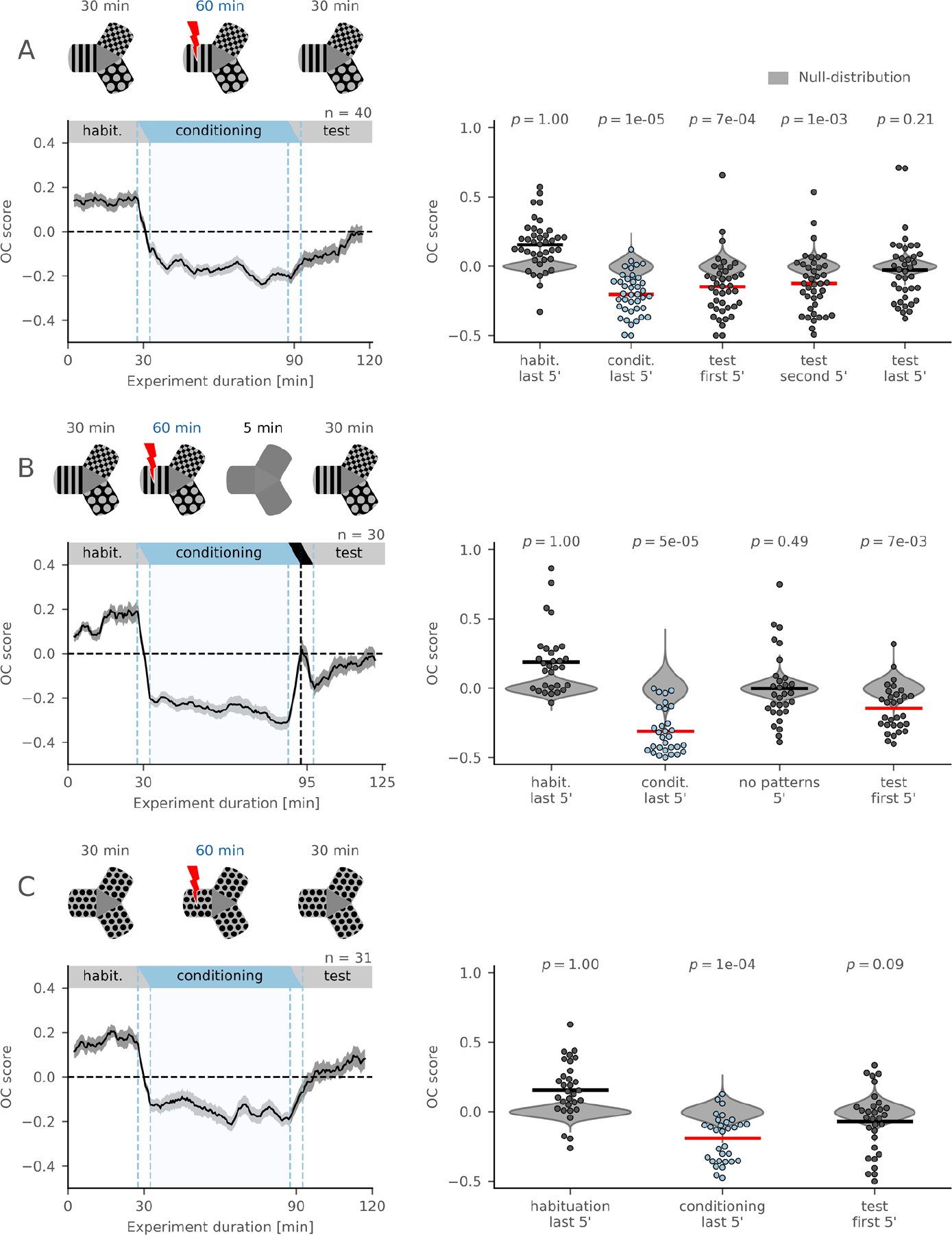
Evaluation of aversive memory formed in the CPA paradigm. **(A)** Top: schematic of the protocol with habituation, conditioning, and test sessions. Bottom, left: OC score moving average. Black line shows average across individuals, gray ribbon shows s.e.m. Bottom, right: comparison of the OC scores in the last 5 min of conditioning and in the first, second, and last 5 min of test session with the null-distribution (permutation test, n = 40 fish). **(B)** Top: schematic of the protocol with habituation, conditioning, no-pattern (all arms with a gray background), and test sessions. Bottom, left: OC score moving average. No-pattern session is indicated on the top of the plot with a black horizontal bar. Note the deflection in the moving average during the no-pattern session (at around minute 95) with a peak value near zero, the chance level. Bottom, right: comparison of the OC scores in the last 5 min of conditioning, 5 min of no-pattern, and in the first 5 min of test session with the null-distribution (permutation test, n = 30 fish). **(C)** Top: schematic of the protocol where visual patterns in all arms are identical. Bottom, left: OC score moving average. Bottom, right: comparison of the OC scores in the last 5 min of conditioning and in the first 5 min of test session with the null-distribution (permutation test, n = 31 fish). All moving averages are calculated with a 5-min time window and a 30-s time step. See also Figure 3 Supplement 1.

To investigate how robust the formed memory was, we introduced a no-pattern session into the experimental protocol between the conditioning and test sessions: after the conditioning session electric stimulation was switched off and all patterns were switched to a gray background for 5 minutes (Figure 3B). During this featureless session, avoidance of the conditioned arm was abolished: the OC score was not significantly different from zero (Figure 3B, permutation test *p*-value = 0.49, n = 30 fish). However, the fish continued to avoid the conditioned arm after the visual patterns were shown again at their original locations (Figure 3B, permutation test *p*-value = 7⋅10^−3^, n = 30 fish). This finding suggests that the fish use visual cues to retrieve an association between a specific place and the aversive stimulus.

To rule out alternative mechanisms, such as odor cues left by the fish after being shocked, we used a Y-maze with three identical patterns (Figure 3C). The occupancy of the conditioned arm was significantly reduced during the conditioning session (Figure 3C, permutation test *p*-value = 10^−4^, n = 31 fish), as predicted by our computational model that takes into account the increased speed induced by electric shocks (Figure 1F, G). However, there was no significant avoidance of the conditioned arm in the test session (Figure 3C, permutation test *p*-value = 0.09, n = 31 fish). This result suggests that fish rely on visual cues, and not on in-maze cues or other sensory modalities, to both retrieve and form an associative memory.

An analysis of EF scores in these three experiments revealed similar effects (Figure 3 Supplement 1). Combined, these results suggest that the memory of the conditioned arm persists for at least 10 minutes after the end of conditioning and that it depends on the presence visual cues.

### Orientation within the electric field predicts strength of response and success of conditioning

We observed a strong variability in individual responses to electric shocks (see the spread of bout amplitudes in Figure 1E, red histogram) and investigated the potential causes behind it. It has previously been reported that fish respond to electric shocks more strongly when oriented parallel to the electric field (Tabor et al., 2014). We confirmed that the probability of response is higher when the fish is aligned with the electric field (Figure 4A, B). Moreover, the strength of the response is significantly higher when the fish is oriented towards the cathode than towards the anode (Figure 4C).

**Figure 4.**
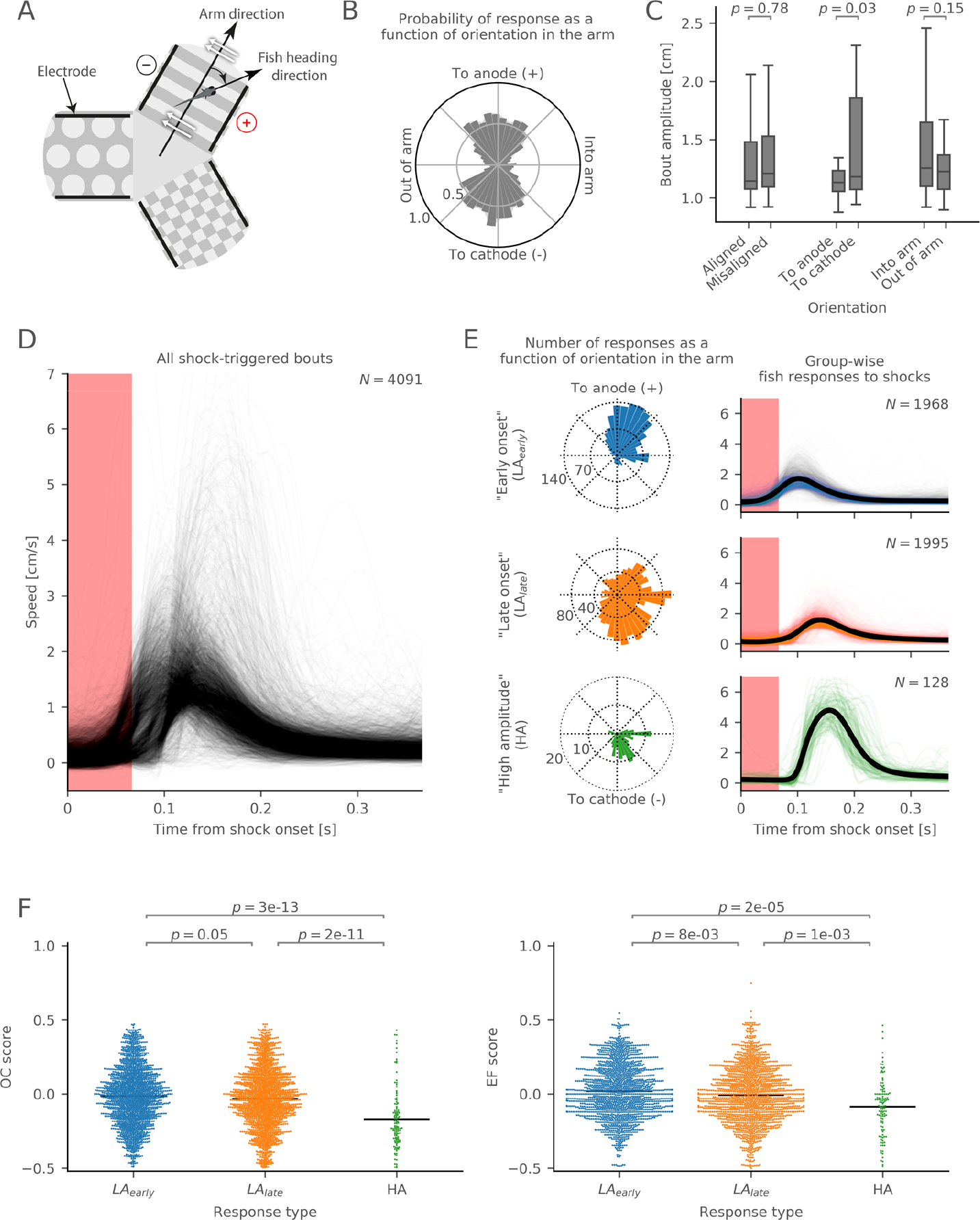
Responsiveness to electric shocks. **(A)** Schematic of a fish’s orientation in the electric field. The orientation angle is calculated between arm direction and fish heading direction. White arrows show the direction of the electric field. Black lines indicate the positions of the electrodes. **(B)** A radial histogram of the probabilities of shock-triggered swim bouts (i.e. responses to shocks) plotted against the fish’s orientation in the electric field (bin size 9°). **(C)** Comparison of the bout amplitudes between different orientations in the electric field (2-sample *t*-test). Bout amplitude is calculated as speed integrated over the duration of a bout. “Aligned” bouts include anode- and cathode-facing orientations; “misaligned” bouts include into-arm- and out-of-arm-facing orientations. **(D)** Variety of amplitudes and onsets of individual shock-triggered swim bouts. Each curve shows the speed during an individual bout (N = 4,091 bouts). **(E)** Response types identified by hierarchical cluster analysis: low amplitude with early onset (LA_early_, N = 1,968 bouts), low amplitude with late onset (LA_late_, N = 1,995 bouts), and high amplitude responses (HA, N = 128 bouts). Left: radial histogram of how many bouts of a certain type occur plotted against the fish’s orientation in the electric field (bin size 9°). Right: speed over time after the shock onset. Y-axes are the same as in (D). **(F)** Comparison of OC (left) and EF (right) scores in a 5-min time window after a shock-triggered bout between different response types (Mann-Whitney test). Each dot represents an OC or EF score after an individual shock-triggered bout. Bouts were obtained from the experiments with 27 fish.

Next, we plotted all individual responses to electric shocks as speed curves in one graph (Figure 4D). We found structure in the amplitudes and onset times of the responses.

Hierarchical clustering with Ward’s linkage method revealed three distinct clusters of response types (Figure 4E). The two low-amplitude (LA) response types differed primarily in their onset time. Moreover, they occurred when fish were in opposing orientations: early onset LA responses occurred mostly in the anode-facing orientation; late onset LA responses tended to occur in the cathode-facing orientation. The third group, high amplitude (HA), responses occurred predominantly when fish were in the cathode-facing orientation.

We hypothesized that different response types correlate with the perceived strength of the shocks, and thus with avoidance level of the conditioned arm. Avoidance level could be quantified by calculating the OC and EF scores in the 5-minute interval immediately after the occurrence of each individual shock response, which we grouped by the response type. The OC and EF scores turned out to be significantly lower after the HA response type than after either early or late onset LA responses (Figure 4F). Thus, the occurrence of HA responses correlated with stronger avoidance of the conditioned arm.

### Spatial learning strategies differ among individual fish

To investigate how avoidance strategies are implemented by individual fish, we grouped their trajectories in the maze in the last 5 minutes of conditioning (see Methods). Each individual trajectory could be described by the occupancies of the three arms and the center. We visualized these occupancies of each animal using a color code. Occupancy of the maze center was coded in shades of gray, where white corresponded to zero occupancy of the center, and black to its exclusive occupancy (zero occupancy of maze arms). Avoidance/preference of the maze arms was coded in a blue-white-red color map, where blue indicates avoidance and red indicates preference of an arm; white color indicates indifference. We used hierarchical clustering on the occupancies and identified three main groups of trajectories: fish preferring the central compartment of the maze to all three arms (4 fish), fish that did not avoid the conditioned arm (9 fish), and fish that avoided the conditioned arm by preferring to stay in one or two safe arms (27 fish) (Figure 5A).

**Figure 5.**
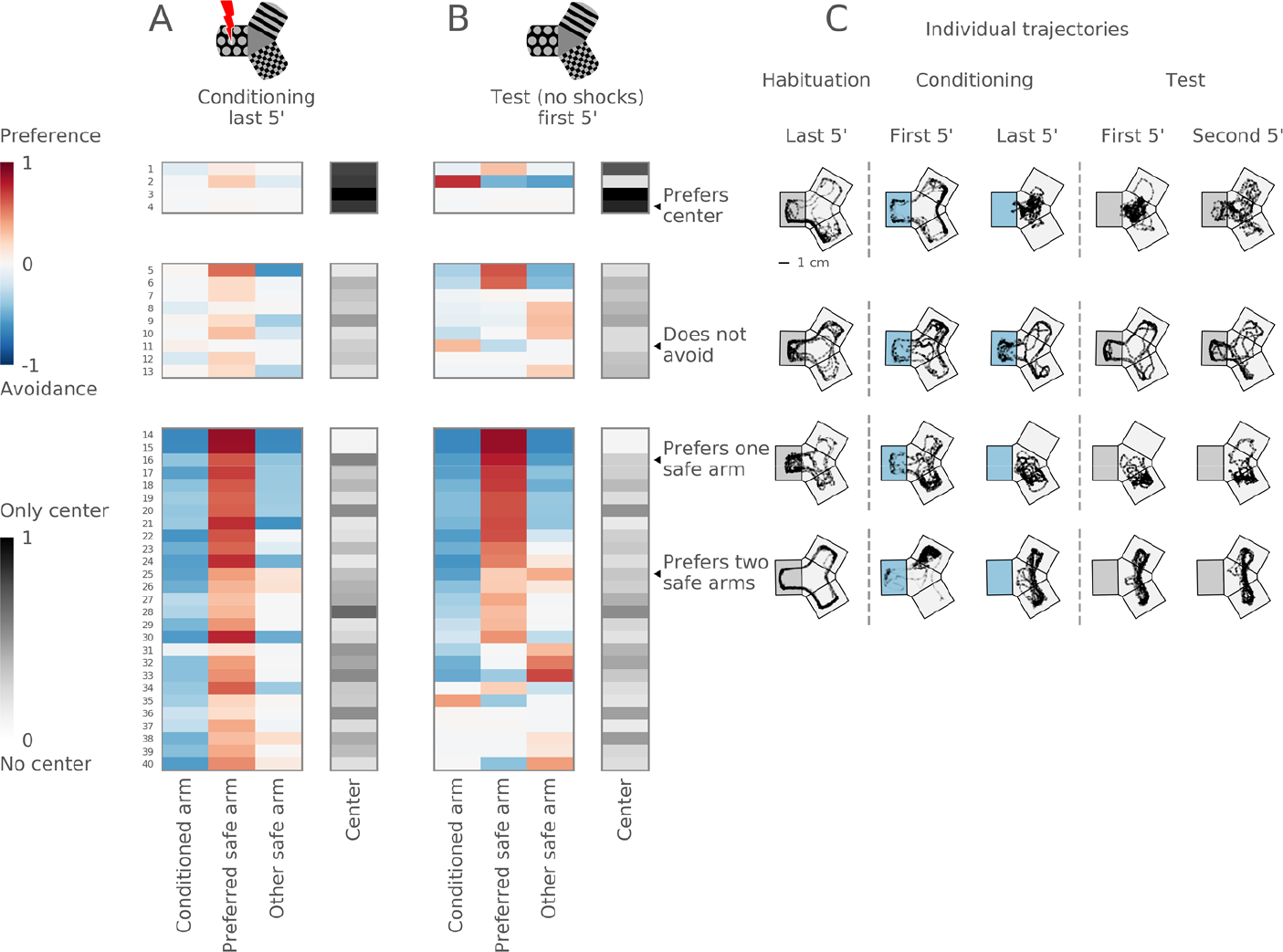
Diverse strategies used to avoid the conditioned arm. **(A)** Top: schematic of the maze during the conditioning session. Bottom: hierarchical clustering of arm occupancies in the last 5 min of the conditioning session reveals three groups: 4 fish preferring the center, 9 fish not avoiding the conditioned arm, and 27 avoiding fish. Each group is presented as a table, one row per fish. Columns correspond to maze arms (from left to right: the conditioned arm, the preferred safe arm, the other safe arm) and the center. Each cell shows a color-coded occupancy value in the particular compartment of the maze for a particular fish, with the logarithmic blue-to-red color scheme for the maze arms and the gray color scheme for the maze center. **(B)** Top: schematic of the maze during the test session. The rows in (A) and (B) correspond to the same fish. Rows within each group are ordered by their similarity to each other in the hierarchical tree in the test session. **(C)** Example trajectories of individual fish in the last 5 min of habituation, in the first and last 5 min of conditioning, and in the first and second 5 min of test session. Top: a fish uses the central compartment as a ‘safe haven’. Upper middle: a non-avoiding fish revisits the conditioned arm despite continued shocks. Lower middle: a fish prefers one safe arm. Bottom: a fish swims in both safe arms. The conditioned arm is depicted with gray background for the habituation session, blue for the conditioning session, and again gray for the test session. The orientation of the conditioned arm varied in the experiments, and is shown here on the left for clarity.

We compared the swimming trajectories of individual fish in the last 5 minutes of conditioning and in the first 5 minutes of the test session, and found that they were stable for the majority of the fish (Figure 5B, see individual examples in 5C). We hypothesized that manipulation of the visual cues could disrupt the stability of swimming patterns in the test session and reveal which cues are relevant for the fish to navigate in the maze.

### Fish may form a “map of safety” of the Y-maze

First, we tested the hypothesis that the fish use a pattern aversion strategy to avoid the conditioned arm. We performed experiments with a new cohort of fish, in which, after conditioning, we replaced the conditioned pattern with a new visual pattern in the same arm (Figure 6A, B, see schematic). Three groups of swimming patterns were observed in this cohort of fish, with center-preferring fish (5 fish), fish not avoiding the conditioned arm (9 fish), and fish avoiding the conditioned arm (20 fish). Interestingly, fish continued to avoid the conditioned arm even after the conditioned pattern was replaced (Figure 6B, blue left column indicates avoidance; see also Figure 6 Supplement 1A-C). Individual trajectories illustrate how fish continue to avoid the conditioned arm (Figure 6C). Control experiments with pattern replacement without conditioning confirmed that replacing one pattern was neither intrinsically aversive nor attractive to the fish (Figure 6 Supplement 2). These results suggest that fish may not only learn to avoid a visual pattern, but also form a “map of safety”, which influences their navigational behavior in the Y-maze.

**Figure 6.**
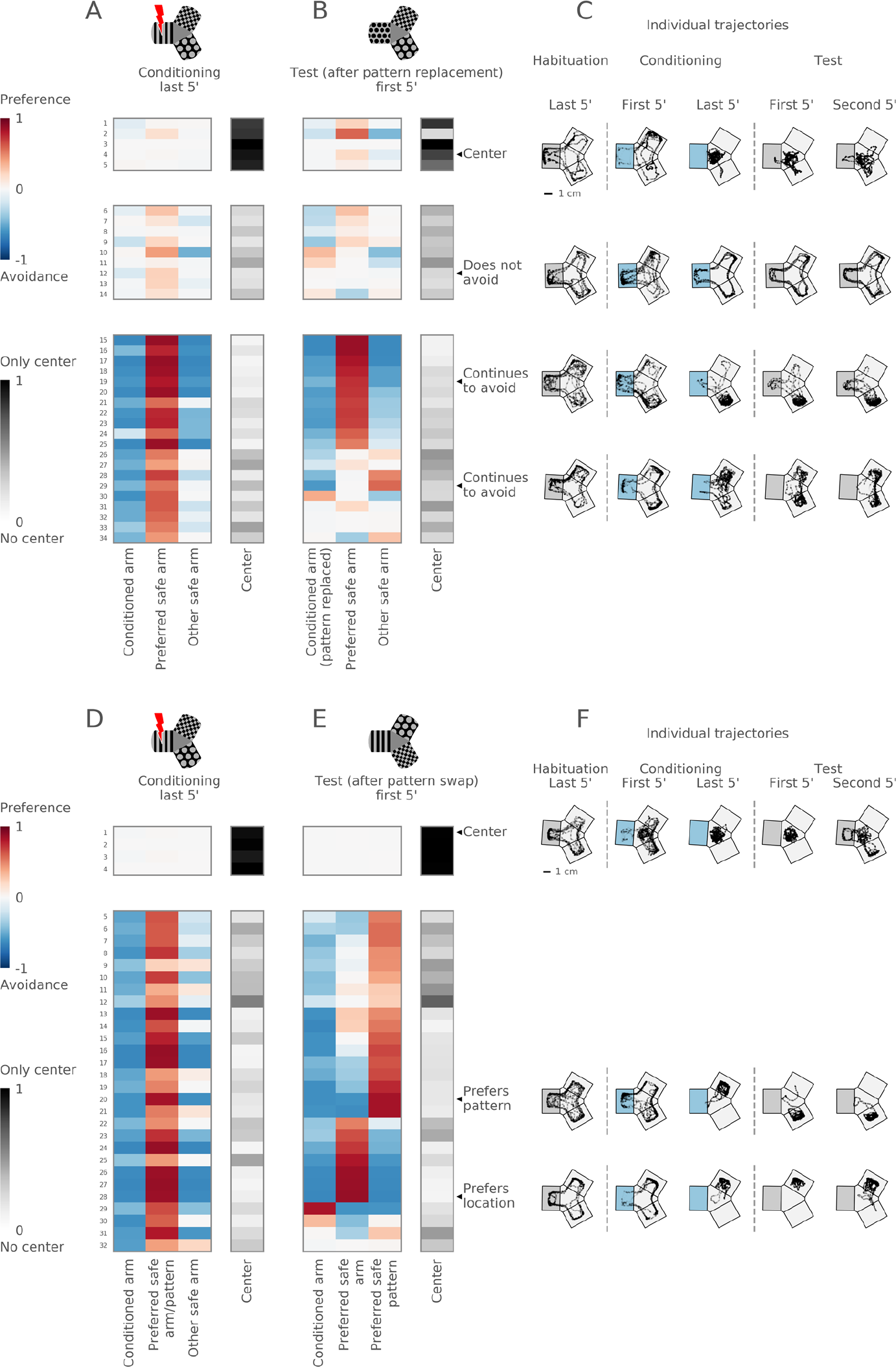
Role of pattern avoidance and preference in the conditioning paradigm. **(A, B)** Top: schematic of the maze during the conditioning and test sessions in experiments with pattern replacement. Bottom: hierarchical clustering reveals three groups: 5 center-preferring, 9 non-avoiding, and 20 avoiding fish. **(C)** Example trajectories of individual fish before and after the pattern replacement. Top: a fish prefers the center. Upper middle: a non-avoiding fish. Lower middle: a fish swims in one safe arm before and after the replacement of the conditioned pattern. Bottom: a fish swims in both safe arms before and after the pattern replacement. **(D, E)** Top: schematic of the maze during the conditioning and test sessions in experiments with pattern swap. Bottom: hierarchical clustering reveals two groups: 4 center-preferring and 28 arm-avoiding fish. **(F)** Example trajectories of individual fish before and after the pattern swap. Top: a fish prefers the center. Middle: a fish prefers one arm and switches the arm after the patterns are swapped. Bottom: another fish also prefers one arm but stays in the same arm after the pattern swap. See also Figure 6 Supplements 1 and 2.

Next, we tested the hypothesis that the fish learn to prefer a safe pattern. We performed experiments in which two safe patterns were swapped after conditioning (Figure 6D, E, see schematic). In these experiments, we identified two groups of center-preferring (4 fish) and safe-arm-preferring individuals (28 fish) (Figure 6D). The analysis of the OC/EF scores showed that the fish successfully avoided the conditioned arm even after the safe patterns were swapped, i.e. the fish retained their aversive memory of the conditioned arm (Figure 6E, blue left column indicates avoidance; see also Figure 6 Supplement 1D-F). After swapping the two safe patterns, we observed two different behaviors. About two thirds of the animals “followed the pattern” and switched their preference to another safe arm, i.e., these fish stayed with the preferred visual pattern (Figure 6E, red in the right column). The other third continued to prefer the same arm despite the pattern swap, i.e., these fish kept the preferred location (Figure 6E, red in the middle column). Individual trajectories illustrate these two different strategies (Figure 6F). This result suggests that fish can use location, rather than rely solely on visual information, in the Y-maze to seek safety.

We next tested the strength of the safety-seeking behavior by creating a conflict between safety and avoidance cues. We rotated the visual patterns after conditioning such that the conditioned pattern was moved into the preferred safe arm. In the test session, we found that some fish continued to swim in their preferred arm despite the presence of the conditioned pattern (Figure 7B, red in the left column; Figure 7C). Other fish transferred their preference according to the pattern rotation, i.e., these fish started to avoid the previously preferred arm (Figure 7B, red in the right column; Figure 7C).

**Figure 7.**
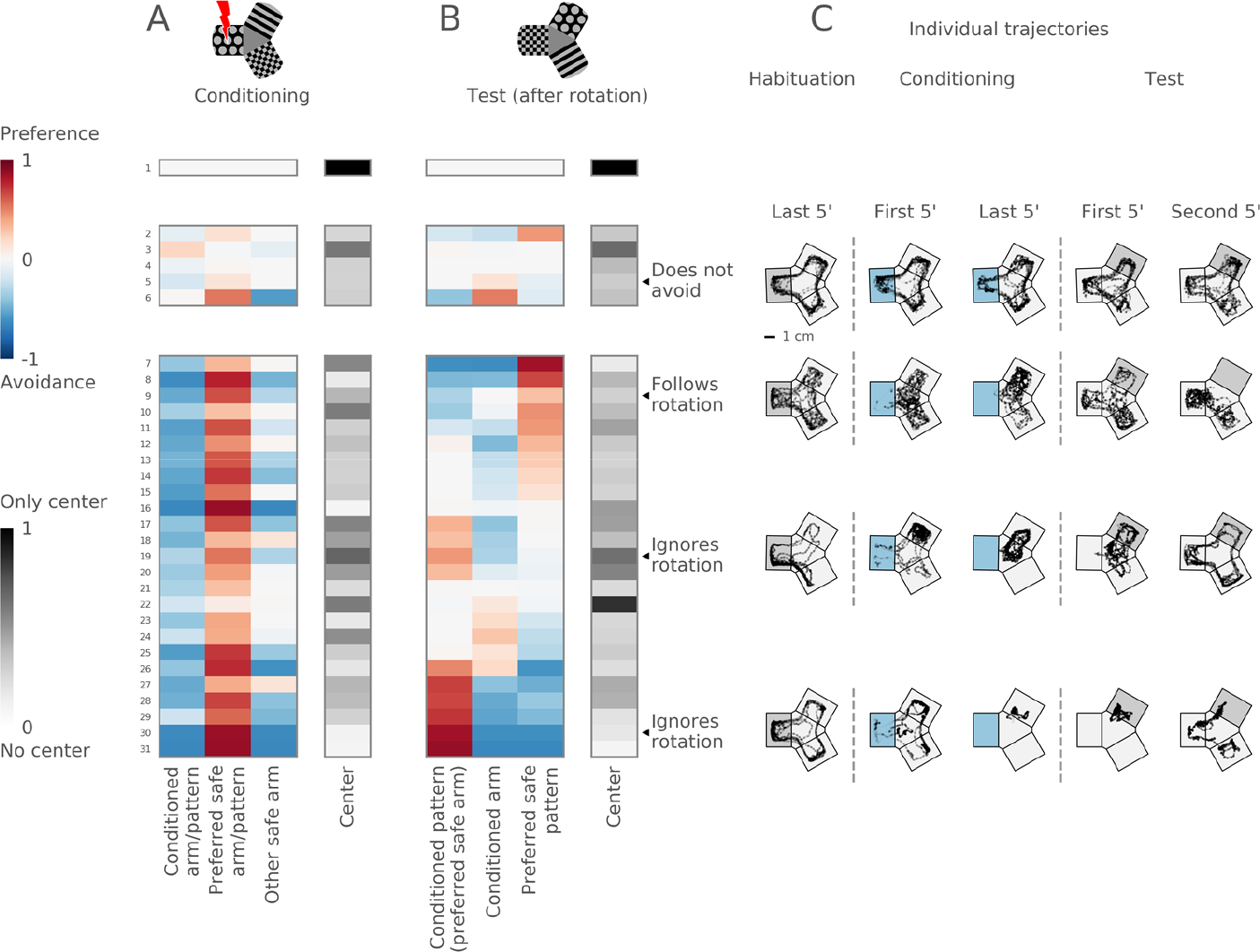
Rotation of visual patterns. (**A, B**) Top: schematic of the maze during the conditioning and test sessions in experiments with pattern rotation. The conditioned pattern moves into the preferred safe arm, thus creating a conflict between avoidance and preference cues. The pattern from the preferred arm moves into the non-preferred arm, pattern from the non-preferred arm moves into the previously conditioned arm. Bottom: hierarchical clustering reveals three groups: 1 center-preferring, 5 non-avoiding and 25 avoiding fish. **(C)** Example trajectories of individual fish before and after the pattern rotation. Top: a non-avoiding fish. Upper middle: a fish moves its preference into the previously conditioned arm, and starts avoiding its previously preferred arm. Lower middle: a fish ignores the rotation and stays in its preferred arm despite the presence of the conditioned pattern. Bottom: another fish also ignores the rotation and stays in its preferred arm.

Overall, we observed that the fish can use different strategies in a spatial learning task, such as learning visual patterns for cue-guided navigation, as well as learning a safe location in the maze, independent of the visual patterns.

## Discussion

To determine whether juvenile zebrafish are capable of spatial learning, we devised conditioned place avoidance (CPA) in a Y-maze as a new behavioral paradigm. In our setup, swimming into one of the three arms is punished with a mild electrical shock, and most of the fish learn to avoid this arm at the end of conditioning. The behavioral chamber is cued with a floor of visual patterns, which can be altered in any desired fashion. The shape of the Y-maze allows for a straightforward readout of which of the four compartments the animal chooses to swim in, i.e., the three arms or the central, uniformly gray compartment. Manipulation of the sensory environment in the maze, via replacement, swapping or rotation of visual patterns, revealed that individual fish use different learning strategies to solve the problem.

Zebrafish older than 3 weeks showed robust responses to conditioning in the CPA paradigm, while larvae at one week post fertilization failed to learn, in agreement with previous observations (Valente et al., 2012). Two-week old zebrafish showed highly variable responses. A possible explanation for this variability is that fish at that age differ strongly in their developmental stage, as is suggested by their widely varying body length (see Figure 2). If brain development correlates with body length, then fish siblings may show different learning capacities depending on their size. This observation can be used in future experiments to select groups of fish based on their developmental maturity rather than chronological age.

The effects of conditioning were assessed with two measures: arm occupancy (OC) and arm entry frequency (EF). While these measures are often correlated (the more frequently the fish enters an arm, the higher the occupancy of that arm, at least if all other motion parameters stay fixed), we demonstrated that they can also decouple due to mechanisms unrelated to learning. Modeling results showed that a lowered occupancy of the conditioned arm during conditioning can be explained by the fish’s reaction to electric shocks (i.e., by increased speed of swim bouts in the conditioned arm), with no learning involved. At the same time, the second measure, the entry frequency of the conditioned arm, was not affected by increased swim speed. These results are consistent with the performance of the young larvae, who do not learn in the CPA paradigm: their occupancy measure decreases during training, but their entry frequency is unaltered.

In juveniles, both OC and EF measures of the conditioned arm decreased during conditioning and persisted at a reduced value for at least 10 minutes after cessation of the shocks. The memory was robust to a brief removal of distinct visual cues between the conditioning and test sessions (see Figure 3). However, the avoidance of the shocked arm diminished by the end of the 30-minute-long test session. Such rapid memory extinction is in contrast with fear conditioning studies in rodents (Fanselow, 1990). Usually, one electric shock is sufficient for a rodent to establish a very strong, lasting aversion to a conditioned location. Fish, on the other hand, repeatedly revisited the conditioned arm of the Y-maze. Such behavior can be interpreted as evidence for poor learning. Alternatively, it can be a sign of a different behavioral strategy. Indeed, in the wild, two behavioral drives compete against each other in the fish: the need to avoid danger and the need to forage. Fish live in water, where the positions of objects, predators and prey change very quickly due to water currents and the three-dimensional environment. Such volatility could reward fish that return to a previously dangerous place, given there is a high probability that the predator is gone and that food has appeared in the meantime.

The mechanism behind the gradual loss of conditioned aversion in the test session could be passive, where fish forget, or active, where fish re-learn the safety of the previously conditioned arm. Active re-learning should depend on the presence of the visual cues (given their relevance in the paradigm; see Figure 3C), and should occur when the fish visits the arm with the conditioned pattern and receives no electric shock, thus building a new association of safety with the previously conditioned arm. Passive memory loss should be a function of time, and take place independently of the presence of the visual cues. Further experiments, such as memory re-instatement though re-introduction of electric shocks, are necessary for distinguishing between these two processes.

The responsiveness to shocks depends on the orientation of the fish in the electric field, a phenomenon that has been observed previously (Tabor et al., 2014). As a consequence, the electric shocks frequently do not elicit a response when the fish is oriented perpendicular to the electric field. This suggests that such shocks might be ineffective as aversive stimuli. To improve the effectiveness of conditioning, the experiment could be amended in at least two ways. Electric shocks could be applied only when the fish is oriented towards the electrodes (i.e. is aligned with the electric field). Alternatively, more electrodes could be added to the setup so that the fish is always facing them, independent of its orientation.

Analysis of individual responses to electric shocks revealed three response types: two low-amplitude types (LA) and one high-amplitude (HA) type. LA response types clearly separated into those correlated with shock onset (early onset) and those correlated with shock offset (late onset). Interestingly, this separation was correlated with the orientation of the fish at the moment of the shock, even though orientation was not used as a classification parameter. In particular, early onset swim bouts occurred when the fish was oriented towards the anode, while late onset swim bouts occurred in fish facing the cathode. We hypothesize that the polarity of the pulse matters for triggering the response. Briefly, two steel-mesh electrodes, such as those used for shock delivery, could act as plates of a capacitor. The capacitor charges during the electric pulse, and discharges after the pulse is switched off, thereby creating a transient current in the opposite direction to the current from the original pulse, hence effectively presenting a pulse in one direction during shock onset and one in the opposite direction during shock offset.

HA swim bouts were less numerous when compared to the LA response types. Previous studies have shown that mild electric pulses directly activate Mauthner cells, bypassing sensory organs (Tabor et al., 2014). Such activation causes the fish to perform a C-bend, resembling the early stage of escape responses – a highly stereotyped tail movement (Liu, Bailey, & Hale, 2012; Temizer, Donovan, Baier, & Semmelhack, 2015). The C-bend resembles the LA response types observed in this study. On the other hand, the HA responses involve a series of powerful swim bouts, and are more variable in dynamics than LA responses. In contrast to LA, HA responses are followed on average by a more aversive response to conditioning (see Figure 4). One could speculate that the LA responses are triggered by direct activation of the Mauthner cell, as previously described, while HA responses are a result of activation of additional circuits in the brain. These could include sensory organs (e.g. lateral line, nociceptors), or additional reticulospinal neurons. Together they might activate brain centers that encode the aversive quality of memories, such as dorsomedial telencephalon, a homologue of the mammalian basolateral amygdala (Mueller, Dong, Berberoglu, & Guo, 2011; Poulos et al., 2009). This observation suggests that electric shocks should be used as aversive stimuli with caution, as some types of observed reactions to shocks (LA) might be artifacts of direct activation of the Mauthner cell and might not have any perceptual or emotional saliency to the animal.

Experiments with identical visual patterns revealed that, on average, the absence of distinct visual patterns prevents the formation of conditioned responses (see Figure 3). This suggests that distinct visual patterns are necessary for learning in the CPA paradigm for the majority of the fish. This, however, does not exclude the possibility that a minority of fish (which only slightly influences the sample average) could use cues other than the projected patterns to avoid the conditioned arm.

Fish responded to conditioning with different strategies. A subgroup of fish stayed in the central compartment of the maze, avoiding all of the arms. This strategy is effectively the ‘safest haven’, and does not require differentiation between the patterns. In this strategy, the fish could use the contrast between patterns in the arms and gray color of the central compartment as a cue for spatial learning; alternatively, the fish could use pattern-independent geometric perception of the central area as a part of the maze most removed from the walls (i.e. the center of the maze is a more open space). Another subgroup consisted of fish preferring one or two safe arms, with the former being more common. A third subgroup consisted of fish that failed to avoid the conditioned arm. This behavior could be explained by an insensitivity of the fish to electric shocks or by a failure to learn.

We hypothesize that these diverse swimming patterns could be a result of associating different external cues with either punishment or safety (Figure 8). One-arm visitors could learn to seek one safe arm, using either its location cue or its visual pattern. Two-arm visitors could learn either to avoid the conditioned arm or to seek two safe arms. Mixed strategies are also plausible. When we replaced the conditioned pattern in the test session, most fish nevertheless continued to avoid the conditioned arm (see Figure 6). Thus, the strategy of using pattern avoidance as a learning cue is not sufficient to explain the avoidance behavior; rather, pattern avoidance is likely combined with a strategy of safety seeking, or avoiding transitions from a safe pattern to a punished location. The swap of two safe patterns after conditioning revealed that two thirds of the fish preferred a safe pattern and switched their arm preference after the pattern swap, suggesting a strategy of learning a safe pattern. On the other hand, the other third preferred a safe location in the maze and stayed in the same arm despite the pattern swap. The existence of location preference was further confirmed in experiments where all three patterns were rotated. There, some fish “rotated” their occupancy preferences together with the visual patterns, while other fish ignored the rotation, even when the conditioned pattern was rotated into the preferred arm of the fish (see Figure 7). Location-preferring fish could combine geometric cues in the maze, such as the corners of an arm, with an egocentric navigation strategy to stay within an arm. Such strategies are, however, prone to error accumulation: without any reference to stable external landmarks animals tend to lose spatial orientation with time. In the case of radially symmetric Y-maze, animals that rely on an egocentric strategy but leave the preferred arm from time to time would rather quickly lose its location; this prediction agrees with the short-term memory of fish observed in our paradigm.

**Figure 8.**
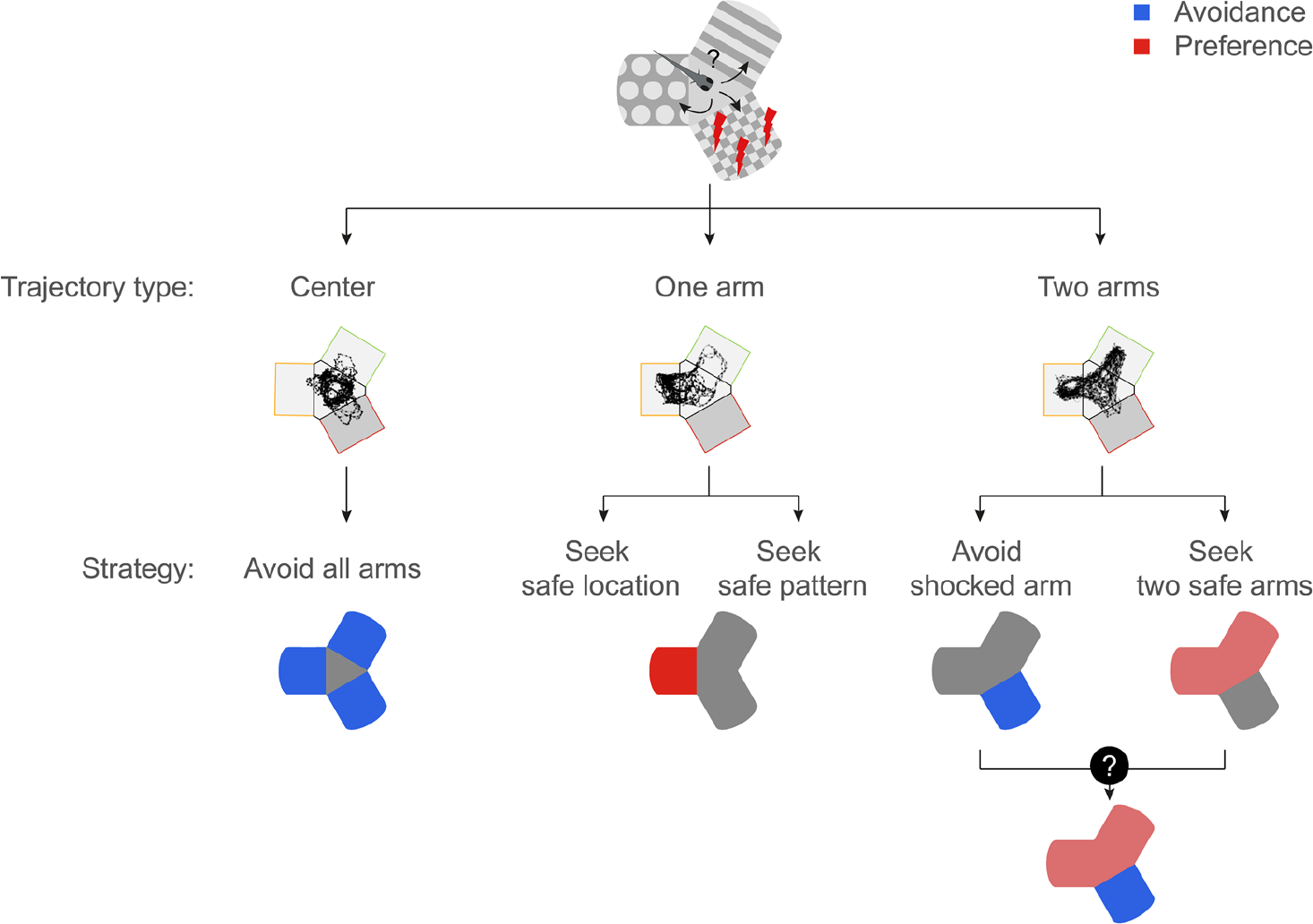
Diverse strategies for conditioned place avoidance among individual animals. Center visitors could avoid all arms. One-arm visitors prefer one arm: either a pattern or a location in the maze. Two-arm visitors avoid the conditioned arm: either by avoiding the conditioned arm or by a more sophisticated strategy, e.g. learning the two safe patterns or the combination of visual and location cues in the maze.

Such broad repertoire of strategies suggests individual flexibility in the spatial representation of safety. Previous studies suggested that each representation could be encoded by a different brain area (Broglio, Rodríguez, Gómez, Arias, & Salas, 2010; O’Keefe & Dostrovsky, 1971; Packard & McGaugh, 1996; Salas et al., 2006). An exciting hypothesis is that these different representations together comprise a cognitive map of the environment, which stores the relationships between various cues and features (Tolman, 1948). Then, different parts of that map can be associated with safety or danger. When two cues come into conflict (e.g. when patterns are swapped or rotated and previously learned visual cues are dissociated from the geometry of the maze), a fish chooses one of the conflicting cues to execute the relevant behavior.

In conclusion, we have described a novel behavioral paradigm for studying spatial learning in juvenile zebrafish. Analysis of the population revealed robust learning effects, which depended on the presence of distinct visual cues. We uncovered several avoidance strategies, which indicate the presence of flexible neural mechanisms underlying the behavior. Recently developed techniques for embedding of juvenile zebrafish (Bergmann et al., 2018; Matsuda et al., 2017; Vendrell-Llopis & Yaksi, 2016), combined with immersive virtual reality setups, make juvenile zebrafish a promising model organism for future studies of brain activity in a navigating animal.

## Materials and methods

Analysis code is available online at https://bitbucket.org/mpinbaierlab/zebramaze/src/master/.

### Fish husbandry

Wild-type zebrafish of strain TL were maintained at 28 degrees on a 14h-10h light-dark cycle. Embryos were obtained by spawning three adult fish pairs simultaneously. Embryos were raised in Danieau’s buffer (17 mM NaCl, 2 mM KCl, 0.12 mM MgSO_2_, 1.8 mM Ca(NO_3_)_2_, 1.5 mM HEPES) for the first 5 days of development. At 6 days post fertilization (dpf) the larvae were transferred to 3.5l tanks with fish system water (approx. 30 animals per tank). From 6 dpf to 20 dpf they were fed twice a day with live Rotifers and dry algae powder (Tetra Aufzuchtsfutter). From day 20 onwards, the diet was smoothly changed to a combination of freshly hatched artemia and Gemma micro dry food (Skretting). Animals were taken out of the fish facility and into the behavior room directly before the start of each experiment. All animal procedures were conducted in accordance with the institutional guidelines of the Max Planck Society and the local government (Regierung von Oberbayern, animal license 55.2-1-54-2532-108-2016).

### Behavioral setup

The setup was custom-built in the lab. The walls and bottom of the Y-maze were laser-cut out of cast acrylic. The maze arms had a 1:1 width-to-length ratio, with a length of 30 mm; the walls were 10 mm high. Each arm opened to the triangular center of the maze. The ends of the arms were rounded. We used two identical mazes in parallel to allow the testing of two fish simultaneously, and therefore increasing the throughput. Walls of the two mazes were opaque to prevent fish from seeing each other during the experiment. Each maze had a piece of diffusive paper underneath for back projection of the visual stimuli. Both mazes and the diffusive paper were placed into a water basin, in order to remove the additional air layer between the screen and the fish and reduce light refraction. The scene was illuminated from below with a custom-built IR LED array to allow behavioral imaging. Both mazes were positioned under a high-speed camera (Ximea USB3.0 model MQ013MG-ON, Münster, Germany), in such a way that the walls did not produce a vertical shadow. The camera had an IR filter to filter out transmitted visible light. The setup was surrounded with black, opaque walls to shield the fish from distracting visual cues in the room.

### Visual stimuli

Visual stimuli, designed in Python, were projected onto the diffusive paper with an LED-projector (LG model PA70G) via a cold mirror. The mirror was positioned at 45° allowing IR light from below to pass through and light from the projector to be reflected onto the diffusive screen. Stimuli were projected under the arms of the maze. The projected stimuli included (a) a pattern with black dots on light gray background; (b) a pattern with light gray dots on black background; (c) a pattern with black and light gray stripes; (d) a checkered pattern of black and light gray colors; (e) uniform gray (RGB = (128, 128, 128)). The light gray color was used instead of white to lower the brightness of the arena as high brightness could increase stress levels of the fish (personal observations, data no shown). The patterns were designed such that the light-to-dark ratio was 1:1 in order to prevent differences in luminance between the stimuli, as larval zebrafish exhibit phototactic behavior (Burgess & Granato, 2007; Orger, Smear, Anstis, & Baier, 2000). The central area always had a uniform gray color (RGB = (135, 135, 135)), which was different from the gray in the arms to ensure that there was a contrast border at the entrance to the arms. Light gray RGB value was (180, 180, 180), black RGB value was (0, 0, 0).

### Electrical stimulation

Each arm contained a pair of electrodes located at the side walls. Each electrode was shaped as a 30-by-10 mm rectangle made out of steel mesh with wire diameter 0.2 mm, aperture 0.5 mm (Mijo Ilic Drahtgewebe-Shop.de, Bergisch Gladbach, Germany) and covered the entire side wall. The electrodes were connected to a constant current stimulator (Digitimer DS3, USA, FL). Electrical stimulation was applied at 1 Hz in the periods between the entry and the exit of the fish from the conditioned arm. Electric pulses lasted between 50 and 100 ms, depending on the experiment (we did not observe differences in responses to shocks with these pulse lengths, data not shown). Pulse amplitude was 0.7 mA (the value was chosen to elicit visible responses to the electric stimulus in all animals, data not shown). The water used in the experiments was obtained from the fish facility (pH 7.5, temperature 28°C, conductivity 650 μS).

### Behavioral tracking

All tracking was performed using custom-written code in Python, including the OpenCV library (Bradski, 2000). Black-and-white images were recorded at 60 fps. The position of the fish was identified in real-time using background subtraction. The background was calculated as a running average of the last 20 seconds of the recording. This time-dependent background was subtracted from the current frame, the result was filtered with a Gaussian filter with a 5×5 pixel kernel to remove point pixel noise, and then binary thresholded. The fish was identified as the contour with the largest area on the thresholded image. Fish position was calculated in real time as the center of mass of the corresponding contour. The identified position was corrected using a Kalman filter to reduce the noise in the recordings (Kalman, 1960). Filter state variable included (x, y) position of the fish and x- and y-projections of the speed, and was updated for every frame of the recording using the observed fish position. Filter model for motion assumed movement with constant speed. Noise along x- and y-coordinates was assumed to be independent.

### Extraction of swimming parameters

Swim bouts were estimated from the time series of fish positions in the maze after the experiment by analyzing the recorded videos. First, speed of the fish was calculated as the Euclidean distance between positions at adjacent time frames. The speed was then filtered using a finite impulse response filter with a low-pass kernel (Parks–McClellan algorithm with 4Hz cutoff frequency) to remove high-frequency noise (Parks & McClellan, 1972). Swim bouts were defined as intervals of the filtered speed curve above a manually set threshold. The swim bout amplitude was calculated by integrating the area under the speed curve between the boundaries of the bout.

The fish’s heading direction was calculated for each frame of the recorded video. First, a contour of the fish was identified in a manner similar to the calculation during the experiment (see Behavioral tracking). The terminal point of the heading vector was at the center of mass of the contour, which roughly corresponded to the head of the fish. The initial point of the vector was at the furthermost point of the contour from the center of mass, which corresponded to the tail tip of the fish. The heading direction was calculated as a relative angle between the heading vector and the horizontal edge of the image.

Fish’s orientation in the arm was calculated as a relative angle between the heading direction of the fish and the orientation of the arm. The orientation of the arm was given by the arm vector, whose initial point was at the maze center, and whose terminal point was at the arm center. Arm orientation was identified as an angle between the arm vector and the horizontal edge of the image.

Fish’s orientations in the arm were divided into four categories: “into the arm” for angles between −45° and 45°; “towards the anode” for angles between 45° and 135°; “out of the arm” for angles between 135° and 225°; “towards the cathode” for angles between 225° and 315°. “Towards the anode” and “towards the cathode” orientations were called aligned with the electric field, while “into the arm” and “out of the arm” orientations were called misaligned with the electric field.

Fish size was calculated as the Euclidean distance between the tip of the head and the tip of the tail, whose positions were manually picked by analyzing recorded videos. To reduce the assessment noise, the length was identified in five randomly picked frames of the video for each fish. Afterwards the final length was obtained by averaging the five handpicked lengths. The accuracy of this procedure was estimated with a coefficient of variation (CV) of fish size, calculated for every fish by dividing the standard deviation of manually measured fish lengths by the mean of those lengths. CVs were calculated for a random sample of all experimentally tested fish (n = 42). The obtained values did not exceed 5% (data not shown).

### Measures for the CPA paradigm

Behavior in the CPA paradigm was assessed with two measures: occupancy of the arms (OC) and entry frequency of the arms (EF). OC of a particular arm, or maze center, was calculated as the proportion of time spent in the respective part of the maze. EF of a particular arm was calculated by dividing the number of times that the fish entered into that arm by the total amount of entries the fish performed into all arms. The two measures could be calculated for the whole time of an experiment as well as for a part of an experiment (in a corresponding time window).

### Preference score

We created a preference score in order to estimate learning effects. The score was calculated for each measure separately as the difference between the mean OC/EF of the conditioned arm and the average of OC/EF of the other two arms (Equation 1 for the occupancy score, *OC*_*score*_, and Equation 2 for the entry frequency score, *EF*_*score*_). Score values were above zero when the conditioned arm was preferred, and below zero when the arm was avoided. Note that the *OC*_*score*_ is calculated from the OC-values of the three arms, irrespective of the OC-value of the maze center.

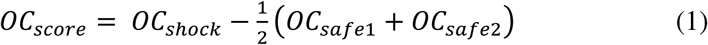

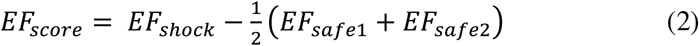

### Moving average

The dynamics of the preference score throughout an experiment were visualized using moving average with a 5-minute time window and a 30-second step, i.e. two adjacent windows had a 4.5-minute overlap. Every point on the moving average represents the average OC/EF score across individual fish in a single time window. Some points of the moving average were calculated from the data of two adjacent sessions (sessions can have different experimental conditions, i.e. ON/OFF electric stimulation); vertical dashed lines mark intervals reflecting mixed experimental conditions.

As individual fish could have different number of arm entries within a 5-minute time window (e.g. because of different swimming speeds), we calculated the weighted average across individual fish for the EF score. The weights were proportional to the total number of entries for a particular fish in a particular time window.

### Experimental protocols

Each experimental protocol consisted of one or more experimental sessions. The sessions followed each other without interruption, and each session could be characterized by its duration, the visual patterns that were projected onto the bottom of the Y-maze, and the ON/OFF status of the electric stimulation. In each protocol, every fish was tested individually and only once.

109 fish that spent longer than one minute in the conditioned arm of the maze (equivalent to experiencing 60 shock pulses) were excluded from the analysis. This was done to prevent excessive stress for the animals.

15 fish that did not reach a stable OC score at the end of conditioning were excluded from the analysis. The score was considered stable if the last point of the moving average lied within the 95^th^ percentile range of the last 10 points of the moving average.

31 fish that showed freezing behavior as a result of conditioning were excluded from the analysis. Freezing behavior was identified by the 5^th^ percentile of the distribution of total distance moved across all fish.

See Supplementary Table 1 for detailed numbers of the animals used for each protocol.

### Modeling

A pseudo-random walk model was designed to investigate how reactions to shocks, independent of learning, could influence the OC and EF measures of the conditioned arm. In the model, the arms of the Y-maze were reduced to three one-dimensional (1D) linear tracks. Each 1D arm had a length (parameter *L*) and a coordinate axis associated with it, with the arm opening located at 0, and the arm end located at a distance *L* from the origin. The center of the maze was modeled as a separate 1D compartment of length *L*_center_. The simulated agent moved along the arm axis with discrete steps (bouts). Each step had a direction (towards the left or the right boundary of the arm) and a size *S*. In order to choose the step size *S*, we used the experimentally observed distribution of swim bout amplitudes (Figure 1E, gray). The experimental distribution was fitted with a Gamma distribution (equation 3) with shape parameter *k* = 1.958 and scale parameter *θ* = 0.999 mm.

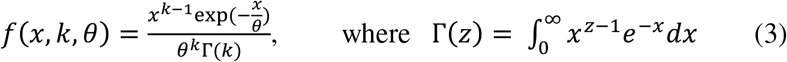

The mean of the fitted distribution, calculated as a product of shape and scale parameters, was equal to 1.956 mm. Thus the average swim bout amplitude, relative to the maze arm length (30 mm), was 1.956 mm/30 mm = 0.065. In order to match these parameters in the model, we drew the step size *S* from a Gamma distribution with mean of 0.065⋅*L*, where *L* was the length of the arm in the modeled maze. Similarly, in terms of *L*, the scale parameter *θ* can be written as *θ* = 0.999 mm ⋅*L*/30 mm = 0.033⋅*L*.

If the simulated agent moved beyond the left arm boundary, it exited its current arm and entered the central compartment. When the simulated agent moved to the right arm boundary, it stopped there until the next step of the simulation (‘sticky’ boundary conditions). Both boundaries of the central compartment were treated equally: if the simulated agent stepped over either the left or the right boundary of the central compartment, it entered an arm. Each arm had a probability of entry associated with it, all probabilities summing to one. The effects of the electric shocks could be simulated in one of the arms. The size of every step made in the shocked arm was multiplied by a parameter *α* ≥ 1, to simulate the increased swimming speed in response to electric shocks.

The model without learning was used to investigate if the increased speed in the conditioned arm alone could explain the changes in OC/EF measures during conditioning. An experiment was simulated with 3 sessions. Each session lasted *n* steps. In the starting habituation session, the step sizes *S* of the simulated agent in all arms were drawn from the same Gamma distribution. At the end of the habituation session, the arm with the highest occupancy was selected as the conditioned arm for the following conditioning session (occupancy was higher in one arm due to stochastic reasons). In the conditioning session, the sampled step size *S* of the simulated agent in the conditioned arm was multiplied by the parameter *α* > 1 to simulate increased speed during the shocks. In the third (test) session, step sizes in all of the arms were again drawn from the same distribution. Probabilities of entry into any of the arms were equal to 1/3 in all sessions. All simulations were run using custom Python code. Parameters used for simulations in Figure 1: *L* = 5; *L*_center_ = 1.5; *k* = 1.96, *θ* = 0.033⋅5 = 0.165; *α* = {1, 2, 3, 4}; *n*_habituation_ = 20,000 steps; *n*_conditioning_ = 40,000 steps; *n*_test_ = 20,000 steps.

A learning component was added to the model to investigate if the learning measures can be used to detect learning effects. In the model with learning, the probability of entry into the arms of the simulated maze was changed from a constant to a variable parameter. Under the learning rule, each entry into the conditioned arm during conditioning decreased the probability of returning to that arm. This rule corresponded to an exponential decay of the probability of entry, with a learning rate *β* and a floor of 0.1 (a non-zero value was chosen because in the experiments the probability of entry never reduced to 0, Equation 4). The probabilities of entry into the other two arms increased correspondingly, to keep the sum of all probabilities equal to one (Equations 5 and 6).

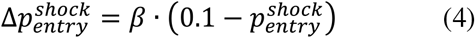

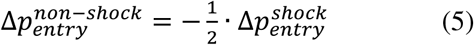

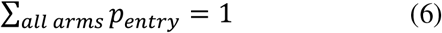

Memory extinction in the test session was simulated by slowly letting the probability of entry into the conditioned arm rise back to the 1/3 level (Equation 7). The update of the probability occurred at every return to the previously conditioned arm. The probabilities of entry into the other two arms were decreased correspondingly (Equation 8 and 9)

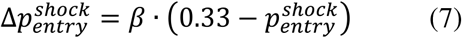

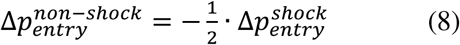

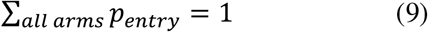

The learning rate *β* used in Figure 1 was equal to 0.03.

### Permutation test

Permutation test was used to assess the differences between the OC/EF measures in the conditioned arm and the other two arms of the maze. For each experimental protocol, a permutation test was performed for the last 5 minutes of the conditioning session (to estimate the effects of shocks during the conditioning) and for the first 5 minutes of the test session (to estimate the memory of the conditioning after the electric stimulation was switched off). OC/EF of each arm for each fish were calculated in these time windows. Then, for each fish separately, the arms were randomly relabeled, so that the OC/EF values were reassigned to different arms. Such relabeling (permutation) was performed n = 10^7^ times. Preference score was then calculated for the experimental values and for each permutation as described above. OC/EF scores obtained from all permutations constitute a distribution of score values for the null hypothesis, i.e. that all arms are interchangeable for the fish, and therefore that there is no significant difference between the OC/EF of the conditioned arm and the other two arms. The experimental score value lies somewhere in this distribution. The significance of the experimental score value is assessed by calculating its position in the null-distribution (Equation 10).

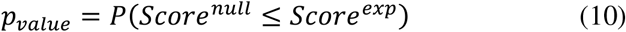

In the cases of very strong effects of conditioning sometimes none of the permutations produced a score lower than the experimental value. In such cases, the estimated *p*-value was equal to 0. In order to get an informative value, we fitted a Gaussian to the null-distribution, and calculated the probability of the experimental value to occur against the Gaussian. This Gaussian estimated *p*-value is reported for the experiments.

### Analysis and clustering of shock-triggered swim bouts

The dataset for shock-triggered swim bouts was obtained from the conditioning sessions of 27 fish. It contained responses to 14,519 shocks, each shock was 67 ms long. For every shock, fish coordinates were extracted for the 35-second interval starting at the shock onset. Every coordinate sequence was then transformed into frame-to-frame speed sequence, calculated from the Euclidean distances between coordinates from adjacent time frames. Every speed curve was smoothed using univariate splines. Speeds whose peak values were lower than a threshold value of 21.4 mm/s were considered non-responses (n = 10,428); the rest were considered swim bouts (n = 4,091). This threshold was chosen such that automatically and visually identified swim bouts matched (data not shown). Additionally, we calculated the orientation of the fish in the conditioned arm at the onset of each identified swim bout.

Principal-component analysis was performed on the dataset of smoothed bout speed curves in order to identify their important features (Jolliffe, 2002). Four principal components could explain more than 90% of total variance in the dataset, and were selected as the main features of the speed curves. Hierarchical clustering with Ward’s linkage method was performed on a bout dataset, which was represented by a 4,091-by-4 matrix (4 principal components per bout, orientation in the arm was not included as a parameter for the clustering). Ward’s linkage method minimizes the sum-of-squares within the clusters (Ward, 1963). The cut level for the cluster tree was chosen so that three clusters emerge. Every cluster was considered to represent a separate type of response to shocks.

Comparison of avoidance levels between different response types was performed using the one-way Kruskal-Wallis test, followed by a post-hoc Mann-Whitney test for group comparisons. Three response types were compared against each other (3 pairs). The avoidance levels were estimated by calculating the OC/EF scores in the next 5 minutes after every individual shock response of a particular type.

### Separation of avoidance strategies

Avoidance strategies were estimated using hierarchical clustering with Ward’s linkage of the arm occupancies in the last 5 minutes of conditioning. For each fish, we computed the occupancy of each arm as described above and re-centered them around zero (so that they sum up to zero). Negative arm occupancy means avoidance, and positive arm occupancy corresponds to preference of the arm. Additionally, we calculated the occupancy of the center. As a result, for each fish a vector of 4 values was calculated: 3 corrected arm occupancies and 1 center occupancy. Importantly, the three occupancies for the arms were linearly dependent, so we excluded one of the arms in clustering. In total, clustering was applied to a 179×3 occupancy matrix, where 179 rows corresponded to individual fish (note that we used fish pooled from different experiments with identical conditioning sessions). Before clustering, values within each column were standardized by centering around the mean and scaling to unit variance.

At each node of the clustering dendrogram, we estimated the significance of splitting the subtree under that node into two clusters. For any current subtree we permuted occupancy values within the columns of the corresponding occupancy matrix (10^4^ permutations in total). We used the same clustering method to get two clusters for each permutation, and obtained linkage values between these two clusters. These linkage values formed a null-distribution corresponding to the hypothesis that there is no separation of the subtree at the investigated node. Then, the position of the original, non-permuted, linkage value was identified in the null-distribution. The significance threshold was chosen to accommodate multiple comparisons between different subtrees of one dendrogram. Briefly, for the node J the significance value 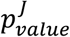 was adjusted by dividing the chosen significance level (0.05) by the number of nodes between the current node and the root of the dendrogram, *D*_*J*_ (Equation 11).

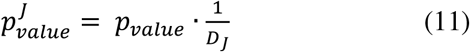

Statistical significance of cluster separation was estimated recursively, starting from the root and investigating each of the following two subtrees. Using this method, we identified six statistically significant clusters. For visualization and explanation purposes, we combined four clusters with the lowest occupancy of the conditioned arm into the ‘arm-avoiding’ group, while the other two clusters corresponded to ‘non-avoiding’ and ‘center-preferring’ groups (see Figure 5).

## Author Contributions

K.Y., A.H. and H.B. designed the study. A.T.C. and K.Y. developed the analysis pipeline and the computational model. K.Y. developed the behavioral assay, carried out the behavioral experiments and performed the data analysis. K.Y., A.H. and H.B. wrote the paper in collaboration with A.T.C.

## Acknowledgments

This work was funded by the Federal Ministry of Education and Research through the Bernstein Center for Computational Neuroscience Munich (01GQ1004A), by the Max Planck Society, and by the German Research Association (DFG) via the RTG 2175 “Perception in Context and its Neural Basis”.

Authors declare no competing interests.

## Supplementary video 1

Fish is swimming in the Y-maze in characteristic bouts. Once it enters the conditioned arm, it receives electric shocks with 1Hz frequency, until it leaves the arm again. Duration of each shock is marked with a red dot. The video is slowed down 4 times. Available at https://web.gin.g-node.org/KseniaYashina/Zebramaze

**Supplementary Table 1.**
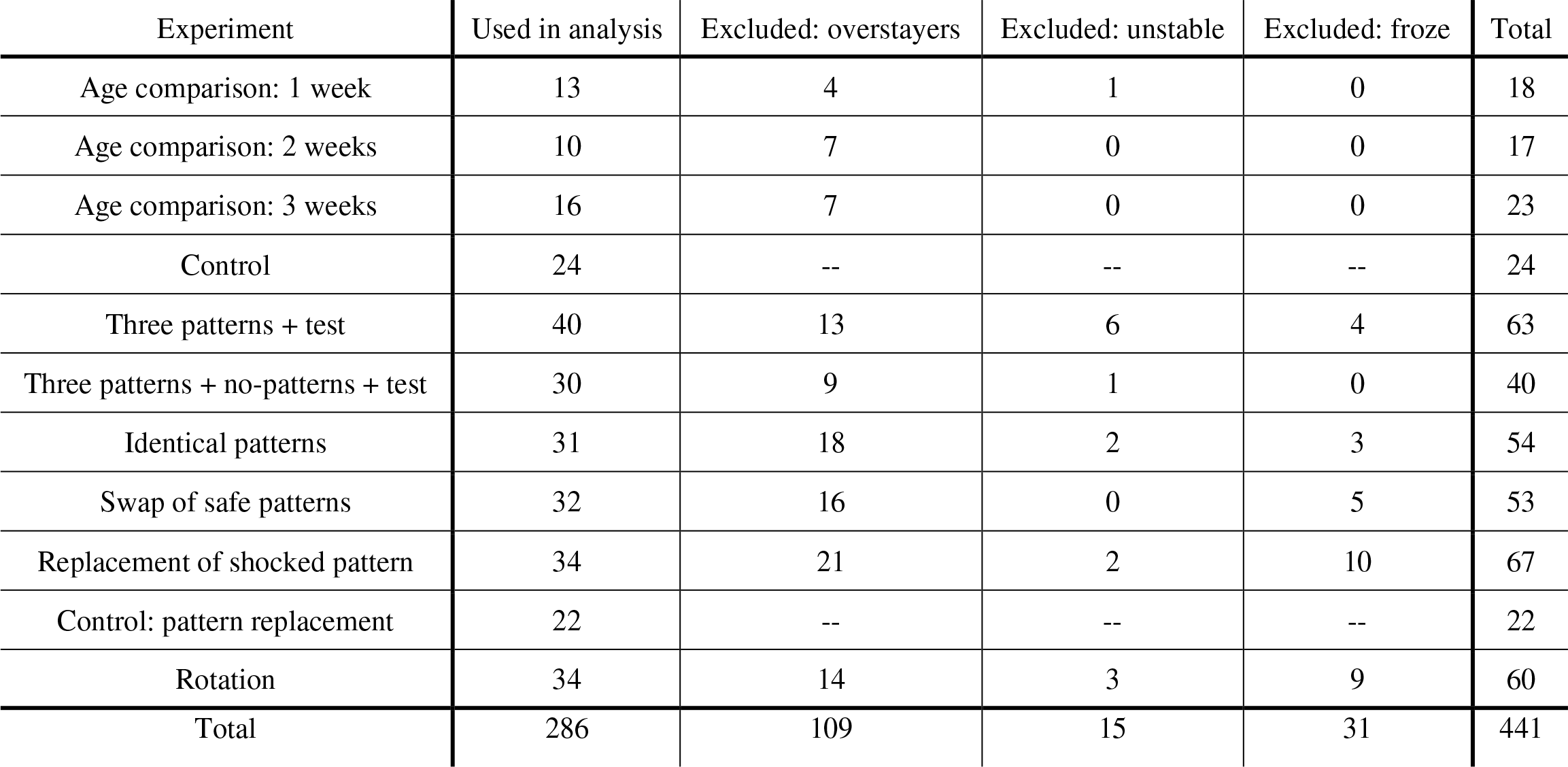
Protocols and animals used in the experiments.

| Experiment | Used in analysis | Excluded: overstayers | Excluded: unstable | Excluded: froze | Total |
| --- | --- | --- | --- | --- | --- |
| Age comparison: 1 week | 13 | 4 | 1 | 0 | 18 |
| Age comparison: 2 weeks | 10 | 7 | 0 | 0 | 17 |
| Age comparison: 3 weeks | 16 | 7 | 0 | 0 | 23 |
| Control | 24 | -- | -- | -- | 24 |
| Three patterns + test | 40 | 13 | 6 | 4 | 63 |
| Three patterns + no-patterns + test | 30 | 9 | 1 | 0 | 40 |
| Identical patterns | 31 | 18 | 2 | 3 | 54 |
| Swap of safe patterns | 32 | 16 | 0 | 5 | 53 |
| Replacement of shocked pattern | 34 | 21 | 2 | 10 | 67 |
| Control: pattern replacement | 22 | -- | -- | -- | 22 |
| Rotation | 34 | 14 | 3 | 9 | 60 |
| Total | 286 | 109 | 15 | 31 | 441 |

**Figure 3 Supplement 1.**
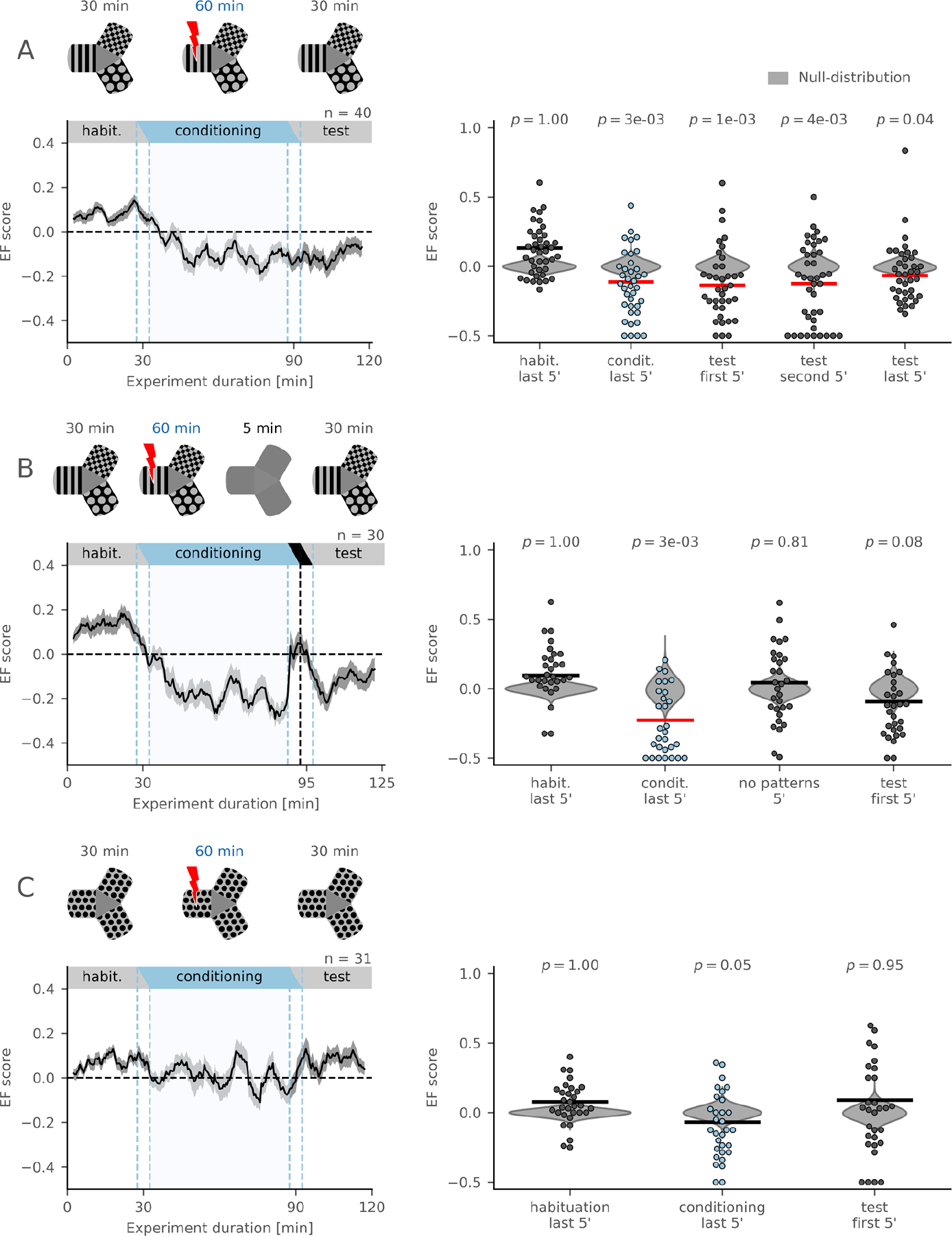
Analysis of the entry frequency score. **(A)** Top: schematic of the protocol with habituation, conditioning, and test sessions. Bottom, left: EF score moving average. Right: comparison of EF scores in the last 5 min of conditioning and the first, second, and last 5 min of test session with the null-distribution (permutation test, n = 40 fish). **(B)** Top: schematic of the protocol with habituation, conditioning, no-pattern, and test sessions. Bottom, left: EF score moving average. Bottom, right: comparison of EF scores in the last 5 min of conditioning, 5 min of no-pattern, and in the first 5 min of test session with the null-distribution (permutation test, n = 30 fish). **(C)** Top: schematic of the protocol where visual patterns in all arms are identical. Bottom, left: EF score moving average. Bottom, right: comparison of EF scores in the last 5 min of conditioning and the first 5 min of test session with the null-distribution (permutation test, n = 31 fish). All annotations are the same as in Figure 3.

**Figure 6 Supplement 1.**
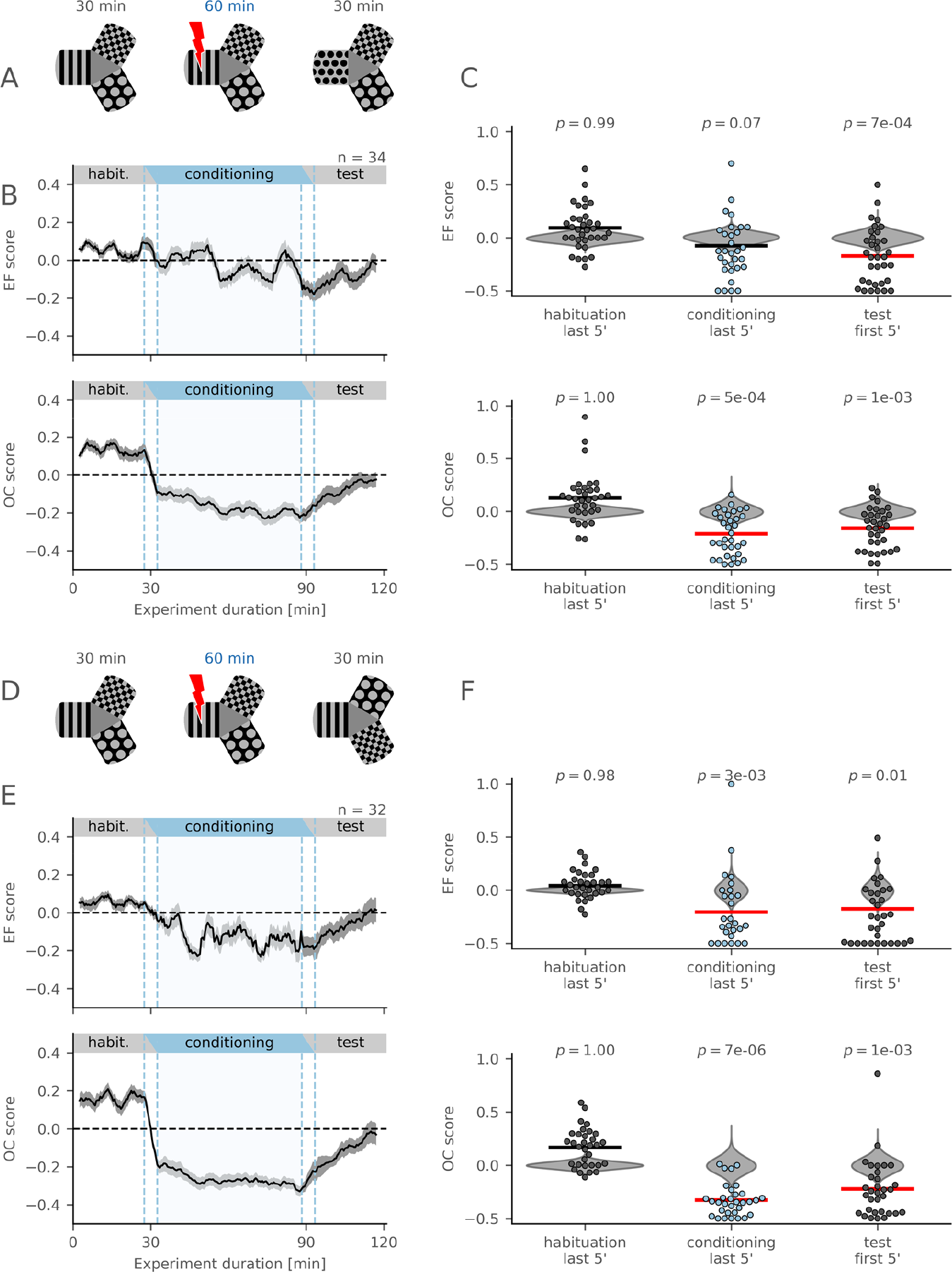
Zebrafish continue to avoid conditioned arm after replacement of conditioned pattern or after swap of safe patterns. **(A)** Schematic of the protocol with the replacement of the conditioned pattern in the test session. **(B)** EF (top) and OC (bottom) score moving averages. **(C)** Comparison of the EF (top) and OC (bottom) scores in the last 5 min of conditioning and in the first 5 min of test session with the null-distribution (permutation test, n = 34 fish). **(D)** Schematic of the protocol with the swap of the safe patterns in the test session. **(E)** EF (top) and OC (bottom) score moving averages. **(F)** Comparison of the EF (top) and OC (bottom) scores in the last 5 min of conditioning and in the first 5 min of the test session with the null-distribution (permutation test, n = 32 fish).

**Figure 6 Supplement 2.**
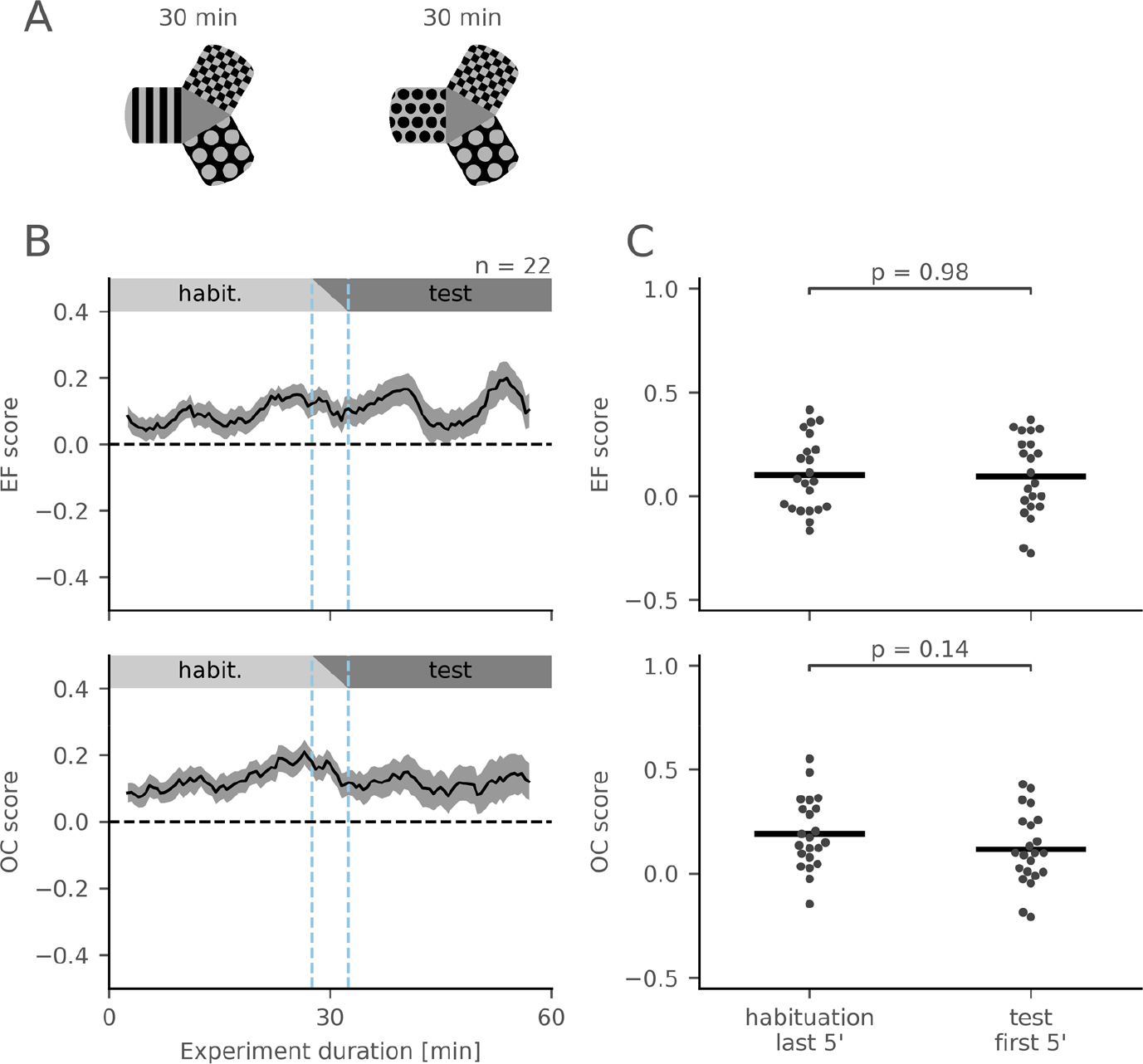
Pattern replacement is neither intrinsically aversive nor attractive to fish. **(A)** Schematic of the control protocol with habituation and test sessions. The pattern in the preferred arm was replaced in the test session. **(B)** EF (top) and OC (bottom) score moving averages. **(C)** Comparison of the EF (top) and OC (bottom) scores between the last 5 min of conditioning and the first 5 min of the test session (two-sided Mann-Whitney test, n = 22 fish). Dots show OC/EF score values of individual fish.

